# Machine learning models to predict *in vivo* drug response via optimal dimensionality reduction of tumour molecular profiles

**DOI:** 10.1101/277772

**Authors:** Linh Nguyen, Stefan Naulaerts, Alexandra Bomane, Alejandra Bruna, Ghita Ghislat, Pedro J. Ballester

## Abstract

Inter-tumour heterogeneity is one of cancer’s most fundamental features. Patient stratification based on drug response prediction is hence needed for effective anti-cancer therapy. However, lessons from the past indicate that single-gene markers of response are rare and/or often fail to achieve a significant impact in clinic. In this context, Machine Learning (ML) is emerging as a particularly promising complementary approach to precision oncology. Here we leverage comprehensive Patient-Derived Xenograft (PDX) pharmacogenomic data sets with dimensionality-reducing ML algorithms with this purpose. Results show that combining multiple gene alterations via ML leads to better discrimination between sensitive and resistant PDXs in 19 of the 26 analysed cases. Highly predictive ML models employing concise gene lists were found for three cases: Paclitaxel (breast cancer), Binimetinib (breast cancer) and Cetuximab (colorectal cancer). Interestingly, each of these ML models identify some responsive PDXs not harbouring the best actionable mutation for that case (such PDXs were missed by those single-gene markers). Moreover, ML multi-gene predictors generally retrieve a much higher proportion of treatment-sensitive PDXs than the corresponding single-gene marker. As PDXs often recapitulate clinical outcomes, these results suggest that many more patients could benefit from precision oncology if multiple ML algorithms were applied to existing clinical pharmacogenomics data, especially those algorithms generating classifiers combining data-selected gene alterations.

## INTRODUCTION

It is now well-established that the efficacy of cancer drugs is strongly patient-dependent. Whereas analgesics such as Cox-2 inhibitors show efficacy in 80% of patients, on average only 25% of oncological patients actually respond to cancer drugs^1^. Consequently, there is a great need to find accurate ways to predict which cancer patients will respond to a given anti-cancer treatment. The predominant approach to date has been to identify a specific somatic mutation to act as single-gene biomarker discriminating between therapy responders and non-responders^2^. Such a predictive biomarker is commonly referred to as an actionable mutation (either a point mutation, deletion or amplication of a specific gene in the tumour sample). Despite being able to predict response to some drugs^3,4^, most patients cannot benefit from single-gene markers because these have not been found for the vast majority of drugs^5,6^. Moreover, drug markers are generally found predictive on a specific cancer type, which means that the marker might not be predictive of drug response on patients from other types^7^.

Not only are these simple drug-gene associations rare, but they are also not strong predictors of drug response in most cases. For example, the mutational status of EGFR in Non-Small Cell Lung Cancer (NSCLC) is a FDA-approved marker of response to Erlotinib^2,8^. The response rate in EGFR-mutant NSCLC tumours was found to be only 16% in this study^8^ (i.e. a 16% precision). Low precision may be due to the interplay of a range of confounding factors, acting on either the same gene (e.g. low expression of mutant EGFR) or other genes (e.g. resistance-inducing mutations in the TP53 gene^9^). The same study unveiled that 67% of the responsive patients were not correctly identified as such, which corresponds to a 33% recall, due to their NSCLC tumours not being EGFR-mutant. This means that two-thirds of NSCLC patients responded to Erlotinib by molecular mechanisms that do not involve EGFR mutations. The Matthews Correlation Coefficient (MCC) summarises both types of error (false positives and false negatives, whose numbers are inversely proportional to precision and recall, respectively) into a single performance metric. This single-gene marker obtained a MCC of just 0.11, which is slightly better than random classification (MCC=0) and very far from perfect classification (MCC=1). Figure 1 clarifies the limitations of this single-gene marker further.

**Figure 1:**
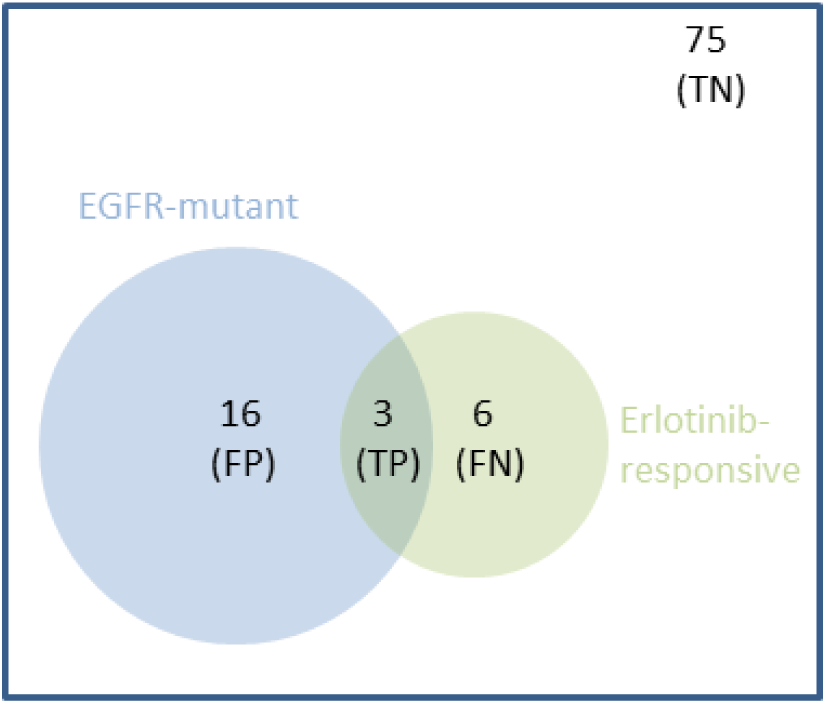
Venn diagram showing the performance of a representative single-gene marker. 100 NSCLC patients were treated with Erlotinib^8^, 19 of them harbouring EGFR-mutant tumours. However, despite being a FDA-approved genomic marker^2^, the mutational status of EGFR was a modest predictor of NSCLC tumour response to Erlotinib. Indeed, 84% (16/19) of EGFR-mutant tumours did not respond. This false positive (FP) rate also means that only 16% (3/19) of these tumours turned out to be responsive (precision of 0.16). Furthermore, 7% (6/81) of NSCLC tumours with wild-type EGFR actually responded to the drug. This false negative (FN) rate means that only 33% (3/9) of the responsive tumours were recalled by this predictor (recall of 0.33). Thus, Erlotinib-NSCLC-EGFR.snv is representative of single-gene markers in that their performance is generally quite modest^5,6^. Intuitively, one can see how combining multiple gene alterations could result in a predictor (blue circle) having a better overlap with the set of responsive tumours (green circle) by decreasing FPs and FNs. The overlap corresponds to the number of true positives (TP). Lastly, patients who do not fall into any of these three non-overlapping categories are true negatives (TN).

While precision and recall vary depending on the drug and its actionable mutation, the example in Figure 1 is representative in that the values of these metrics are generally quite modest^5,6^. Furthermore, it is widely believed that a suitable cancer treatment can only be predicted for those patients who have one of such actionable mutations^5^. We argue however that single-gene markers of drug response constitute only one possible approach to precision oncology. Consequently, more patients could benefit from taking an alternative approach that instead captures the interplay between multiple gene alterations that co-operatively control treatment response within tumours of a specific cancer type.

A promising complement to single-gene markers is the application of Machine Learning (ML)^10^ to learn which combinations of gene alterations are most predictive of *in vivo* tumour response to a given treatment. ML algorithms can build *in silico* models with higher precision (i.e. fewer false positives) by learning which gene alterations, other than the single actionable mutation, influence drug response and how. Regarding increasing recall (i.e. fewer false negatives), ML can potentially learn all the different ways with which tumours of a given cancer type respond to a specific anti-cancer therapy. In that case, ML models would correctly identify not only responders with the actionable mutation as the single-gene marker, but also the responders that are wild-type for that gene. Moreover, ML can provide predictive multi-gene models for some of the many drugs for which a single somatic mutation is simply not enough to predict tumour response^11,12^.

Unfortunately, the limiting factor for the application of ML to this problem is the availability of relevant data. Although the public release of new clinical pharmacogenomics data sets to power precision oncology is often promised, drug response data is typically excluded from these sets or at the very least limited to a few drug treatments. For example, the first release of the AACR Project Genie^13^ contained 19,000 molecularly-profiled tumour samples from various cancer types. However, the responses of the corresponding cancer patients to the administered treatments are still withheld to this date. Even if that information was revealed, cancer patients usually receive drug combinations and several lines of therapy after sample collection, hampering direct associations. This hindrances the discovery of new predictors of drug response. Deep molecular profiling, with no treatment response data, is only part of the puzzle.

In the last six years, data comprising thousands of molecularly-profiled cell lines treated with hundreds of cancer drugs have been made freely available (e.g. GDSC^14^, CCLE^15^ or CTRP^16^). Cell lines are relatively cheap, quick to grow and amenable to high-throughput experiments^17^. In addition, some cell lines have been shown to mimic sufficiently well primary tumours^17–19^. Given their suitability for high-throughput experiments^17^, they are also the model for which more data is publicly available. Thus, such *in vitro* pharmacogenomics data sets are the only ones available to predict response on many pairs drug-cancer type. Despite these advantages, cell lines suffer from several inherent limitations. For example, intra-tumour heterogeneity, extracellular environment and immune system response processes are not captured by cell lines. They are furthermore prone to divergence across passages^18^.

In this context, Patient-Derived Xenograft (PDX) models are of great importance when no relevant clinical data is available^20–23^. Indeed, PDXs effectively capture patient-to-patient response variability to anti-cancer therapy^24,25^. In addition, these preclinical tools faithfully preserve the intra- and inter-tumoral heterogeneity observed in the originating cancer sample and the clinical population, respectively^26,27^. Taken together, these results support the use of these PDX models in guiding clinical therapeutic decisions for a more effective cancer treatment management^28,29^. PDX pharmacogenomics data represents an attractive opportunity to build ML models to predict tumour response in those treatment-cancer type pairs for which clinical data sets are not available. Prominent among such data sets stands the NIBR-PDXE^29^ resource for its high number of PDXs and their comprehensive profiling. Over 1000 PDXs were established, with 40% of these molecularly-profiled at three levels: whole-exome single-nucleotide variants (SNVs), copy-number alterations (CNAs) and gene expression (GEX). Importantly, some of these PDXs were also evaluated with a panel of 60 treatments, which makes these data sets amenable to ML modelling.

Here we carry out a systematic study to investigate how ML can improve the prediction of *in vivo* drug response from tumour molecular profiles by combining multiple gene alterations. To the best of our knowledge, this is the first time that NIBR-PDXE data is analysed with this aim. Note that previous studies applying ML to this problem across multiple drugs have been based on pharmacogenomics data from *in vitro* cell lines, instead of PDXs. There are three additional aspects of this paper that are novel: 1) directly comparing the performance of ML classifiers against that of single-gene markers across multiple treatments, 2) introducing and applying a variant of RF intended to provide a more stringent Feature Selection (FS)^30^ as a way to better handle the high-dimensionality of data sets, and 3) adopting a non-competitive approach where the goal is to identify the most suitable profile and classifier type for each treatment rather than that with the highest average performance across treatments (the current “one-size-fits-all” approach).

Given that single-gene markers based on somatic mutation data are central to current genomic medicine^6,31^, it is surprising that practically all studies evaluating multi-gene ML models have not included a direct comparison with single-gene markers. When we recently carried out such comparison *in vitro* across 127 drugs^12^, we observed that ML models combining multiple gene alterations identified a higher proportion of drug-sensitive cell lines (i.e. had a higher recall) in 93% of the drugs. Here we will determine whether this is also the case *in vivo*. If confirmed *in vivo*, this will mean that many more patients could benefit from precision oncology if multi-gene ML methods are applied to existing clinical pharmacogenomics data.

One major challenge to build any predictive model is the high dimensionality of pharmacogenomics data. Indeed, while typically only tens of tumours have their response to the drug available, the molecular profiles of these tumours may easily aggregate over 50,000 features. To face this challenge, ML algorithms with built-in FS such as Elastic Nets^32–36^, Ridge^33,36^, LASSO^33,34,36,37^ or Random Forest (RF)^12,33,35,36,38,39^ have been used to model pharmacogenomics data from *in vitro* cell lines. For instance, RF ignores those features irrelevant for predicting drug response and thus has been able to tackle to some extent this challenge. However, these methods have also been found to be unable to provide predictive models for many drugs. Here we will employ a new strategy, Optimal Model Complexity (OMC), to complement the ability of RF to reduce the dimensionality of tumour molecular profiles. With OMC, only a very small proportion of the typically thousands of features in the considered molecular profile will be employed by the resulting model. This is beneficial in that much fewer features would have to be experimentally determined in forthcoming tumours. In addition, in those cases where a few tumour features control drug response, predictions should tend to be more accurate because RF will no longer be considering the thousands of irrelevant features that promote model overfitting.

Lastly, the application of a ML method achieving slightly better performance than a previous method on average across drugs is commonly reported. However, the performance of ML methods with similar average performance across drugs can be very different on a particular drug. We therefore adopt a non-competitive approach instead, where the goal is to identify the most suitable profile and classifier type for each treatment. This approach should be collectively more predictive than using the method with the highest average performance across treatments. Another expected advantage is that we will be able to find out which molecular profile is most predictive for a given drug, which is valuable to reduce the time and cost that would be associated to determining the rest of profiles on patients treated with that drug.

## RESULTS

We started the analysis by determining which drug-cancer type pairs in NIBR-PDXE^29^ have sufficient data to be likely to lead to predictive models (see the Methods section). We identified 13 of such treatments in Breast Cancer (BRCA) and another set of 13 treatments in Colorectal Cancer (CRC). All but one of these 26 treatment-cancer type pairs had at least 35 PDXs, each PDX with treatment-response, SNV, CNA and GEX profiles.

### Establishing the best multi-gene predictor for each treatment and cancer type pair

To perform this task, we trained and evaluated two ML algorithms on each data set using leave-one-out cross-validation (LOOCV) as detailed in the Methods section. The first algorithm is RF using all available features (RF-all), whereas the second is an OMC variant of RF to identify the most predictive features in each case (RF-OMC). To account for their stochastic nature, we trained each algorithm on each LOOCV training fold 10 times, thus obtaining 10 estimations of each performance metric per case. Note that this study always reports the median performances of each algorithm on held-out PDXs not used to train or select the model providing the prediction. For instance, each reported MCC is the median of 10 MCC determinations from 10 independent nested LOOCVs. To assess which cases are better predicted by a particular gene, we also performed the standard single-gene analysis to evaluate which genes sensitise PDXs to the treatment when an actionable SNV is detected. In particular, we identified the sensitive marker with the lowest p-value and reported its LOOCV performance. Full results can be found in the single-gene_markers tab of the Supplementary Tables.

Figure 2 shows the results for each of the 26 cases. We found out that the accuracy in predicting treatment response on left-out PDXs strongly depends on the considered treatment, molecular profile and classifier type in both cancer types. The performances of the best predictors from each case range from being slightly above purely random classification (MCC=0) to high in the context of this problem (MCC=0.57). In addition, a large variability is often obtained across the four molecular profiles within the ML algorithm. For each ML algorithm, we show the performance of the most predictive molecular profile for that case. As CNA is just a binarisation of the real-valued copy number (CN) profile, it is not surprising that CN is used much more often than CNA in the best models across cases (19 vs 3 models in Figure 2, respectively). Performance also varies strongly across the three model types within a given case.

**Figure 2:**
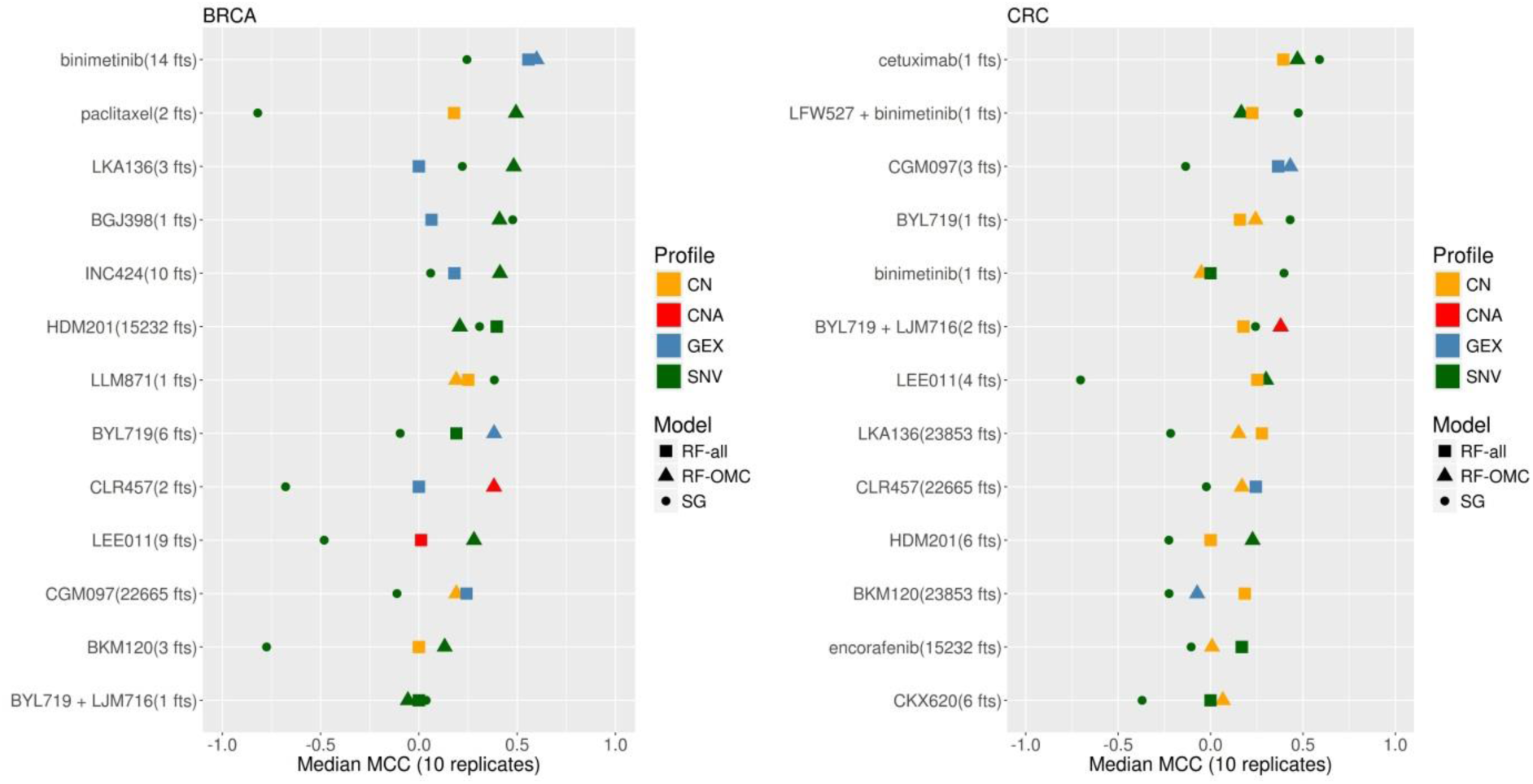
Predictive performance of the best single-gene (SG) marker, Random Forest (RF) with Optimal Model Complexity (RF-OMC) and RF using all features (RF-all). (left) Best predictor per treatment and classification model type on BRCA PDXs. Each row shows the results for one treatment and shows the number of top features with which the best classifier for that treatment was trained on. Furthermore, the colour and shape of each classifier indicates the employed molecular profile and model type, respectively. For instance, paclitaxel (2 fts) appears as a green triangle, which means that Paclitaxel had RF-OMC-SNV as the classifier with the largest median MCC on LOOCV held-out PDXs and this classifier employed the top 2 features from the SNV profile (MUC20 and UPK3BL). All treatments have at least one predictor performing above random level (MCC=0), with the accuracy in predicting BRCA PDX response being strongly treatment-dependent. Importantly for clinical implementation, the best classifier per treatment is usually a model that only requires a handful of features to operate. **(right)** Best predictor per treatment and classifier type on CRC PDXs. Strong predictors of treatment response were also found for this cancer type. However, there were fewer of these predictive models in CRC than in BRCA (five treatments were predicted with MCC>0.4 in BRCA, but only two treatments were predicted at this level in CRC). Top models are more frequently associated with CN profiles in CRC than in BRCA. It is also clear that CNA profiles, using CN as a binary feature (altered/wild type), leads to less predictive models than real-valued CN.

RF-OMC not only leads to more accurate predictors in 14 of the 26 cases, but these predictors merely require a very small subset of all gene alterations to operate (these concise gene lists are reported in tab RF_predictors of Supplementary Tables). By contrast, solely 5 of the 26 cases were better predicted by RF-all. Figure 2 shows that the MCCs of RF-OMC across the 13 treatments were not better than those of RF-all in CRC (P=0.47 from a one-sided paired t-test, both algorithms obtaining an average MCC of 0.19). However, RF-OMC was found to outperform RF-all in BRCA (P=0.009 from a one-sided paired t-test, average MCCs of 0.32 and 0.16, respectively). The best predictor in the remaining 7 cases was a single-gene marker. Overall, these results stress the importance of considering several model types and profiles to predict *in vivo* treatment response.

We have also compared each of these two ML classifiers against a random model where response class is predicted using the prior probability of a PDX to be sensitive. Supplementary figures 2 and 3 show that RF-OMC predicts 21 of the 26 cases better than random (P<0.05; one-sided paired t-test). By contrast, supplementary figures 4 and 5 show that RF-all is only able to predict 16 of the 26 cases at this level (P<0.05; one-sided paired t-test).

**Figure 3:**
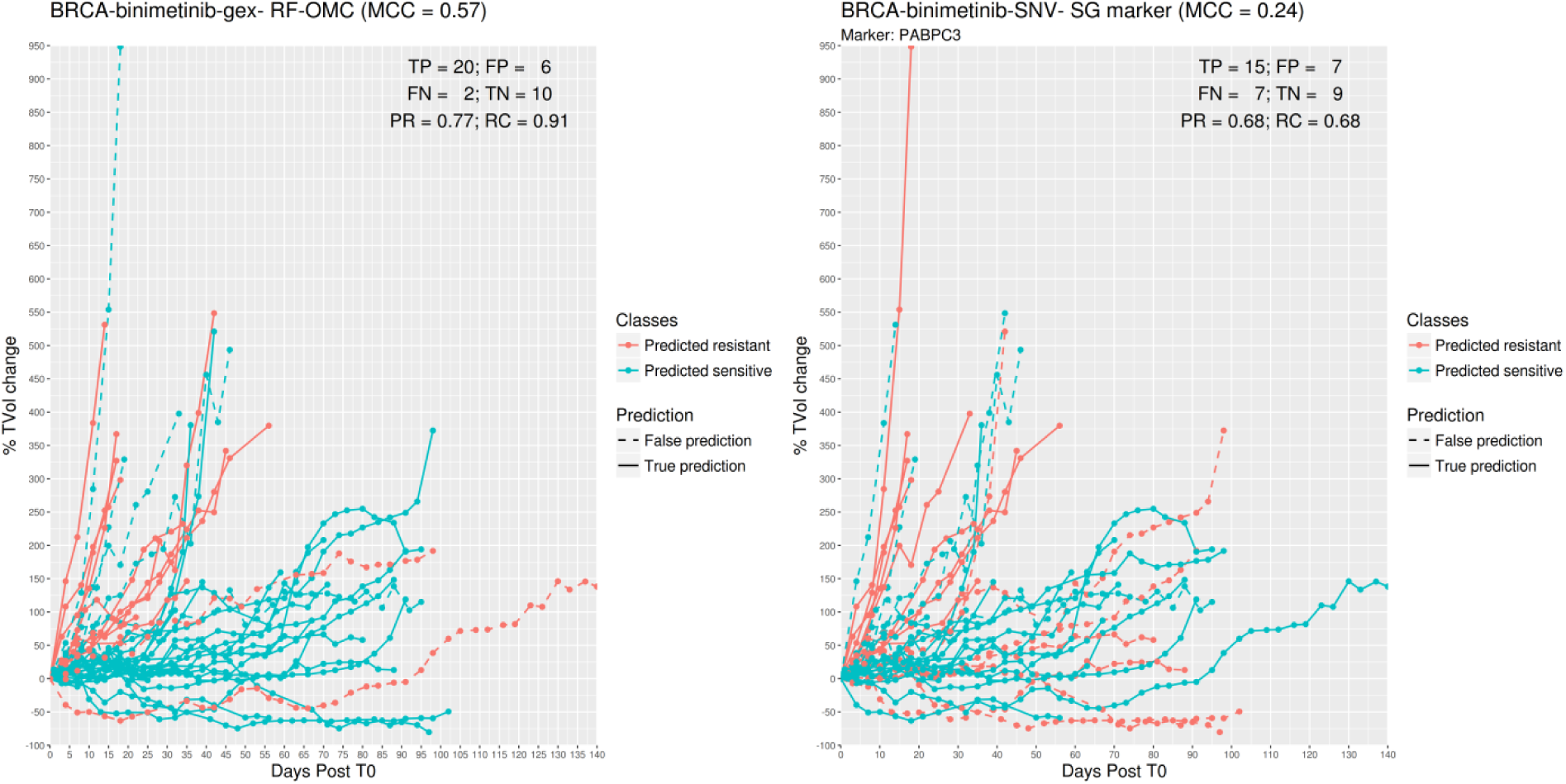
Predicting BRCA PDX response to the MAPK inhibitor Binimetinib. Each line represents one PDX. The vertical axis shows the % of tumour change relative to the tumour volume at time zero (T0; i.e. immediately before the first administration of the drug) and the horizontal axis shows when measurement time in days after T0. From the legend, red discontinuous lines represent false negatives (responsive PDXs that were predicted to be non-responsive) and blue discontinuous lines represent false positives (non-responsive PDXs that were predicted responsive). The better the predictor is, the higher the proportions of lines at the bottom appear in blue (higher recall) and at the top in red (higher precision). **(left)** *Binimetinib (14 fts)*: RF-OMC model predicts BRCA tumour response to Binimetinib by optimally combining the expression levels of only 14 genes (CRB3, NDUFA1, MPG, ECI1, ING2, KIF9, TSTD1, FAM100A, TCEAL3, HAGH, PEX11G, SNORA72, SNORA70 and PIN1). A high level of predictive accuracy was achieved on PDXs not used to train the model: MCC=0.57 (PR=0.77 and RC=0.91). **(right)** *Binimetinib (1 fts)*: Using the same input data and evaluation protocol, the best single-gene marker of Binimetinib sensitivity was the mutational state of PABPC3. The RF-OMC model obtained a substantially higher level of prediction than that of this standard single-gene procedure: MCC=0.24 (PR=0.68 and RC=0.68).

**Figure 4:**
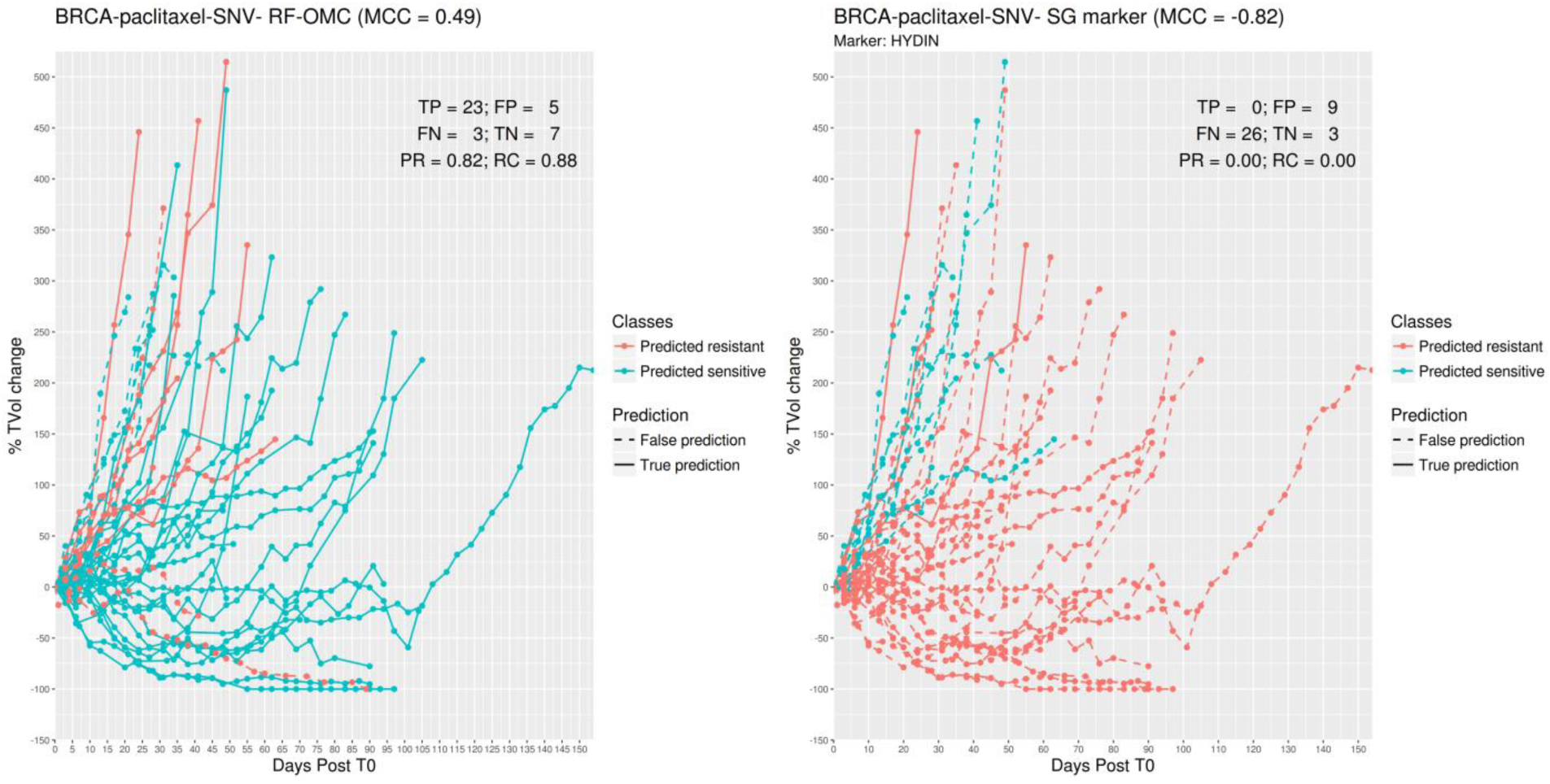
Predicting BRCA PDX response to the tubulin inhibitor Paclitaxel. **(left)** *Paclitaxel (2 fts)*: RF-OMC predicts BRCA tumour response to Paclitaxel by optimally combining the mutational states of 2 genes (MUC20 and UPK3BL). A high level of predictive accuracy was achieved on PDXs not used to train the model: MCC=0.49 (PR=0.82 and RC=0.88). **(right)** *Paclitaxel (1 fts)*: Using the same input data and evaluation protocol, the best single-gene marker of Paclitaxel sensitivity was the mutational state of HYDIN. The RF-OMC model obtained a much higher level of prediction than that of this standard single-gene procedure: MCC=-0.82 (PR=0.00 and RC=0.00).

**Figure 5:**
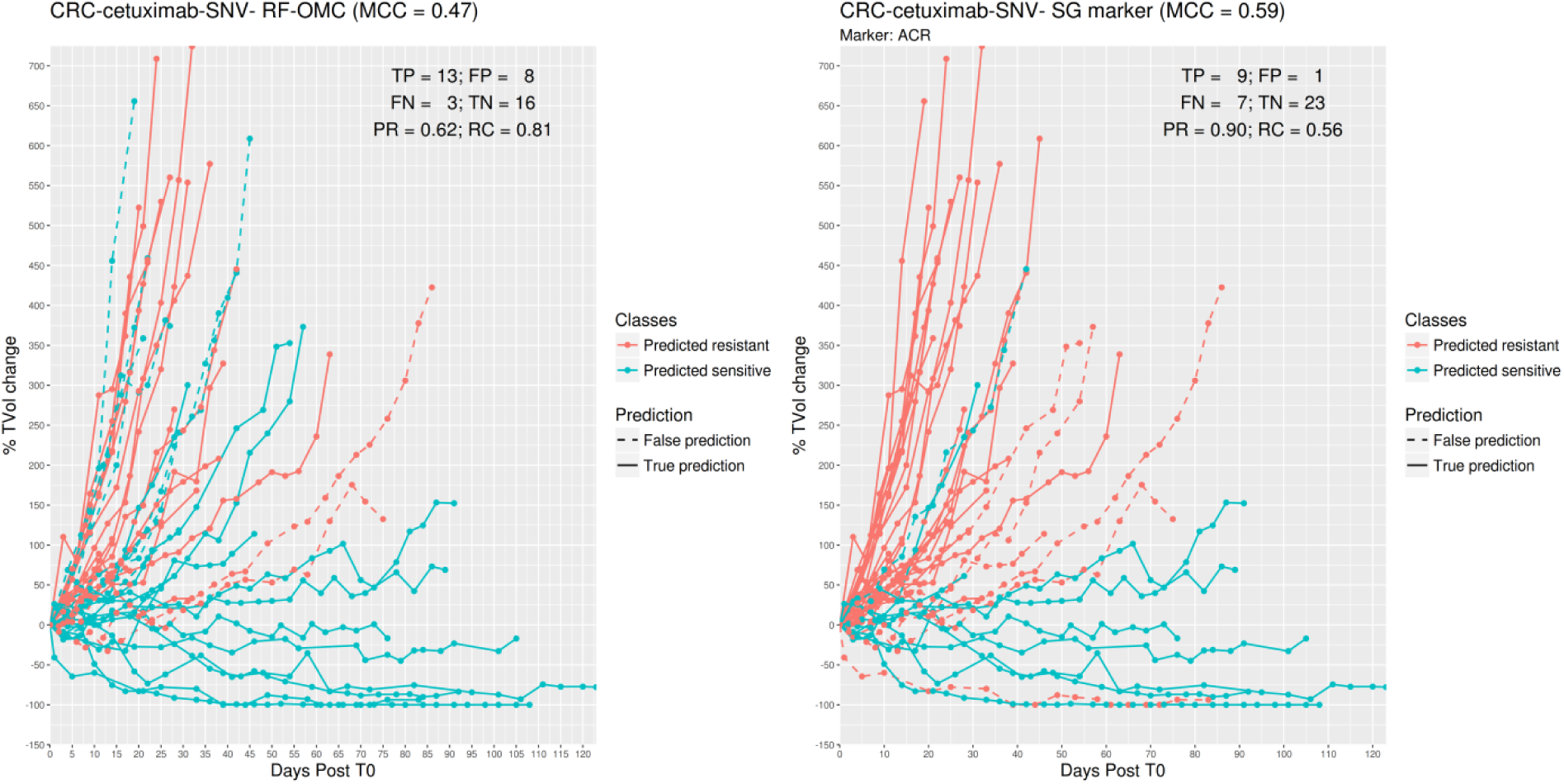
Predicting CRC PDX response to the EGFR inhibitor Cetuximab. **(left)** *Cetuximab (4 fts)*: RF-OMC predicts CRC tumour response to Cetuximab by optimally combining the mutational states of 4 genes (ACR, DENND4B, NOTCH1 and RPL22). Again, a high level of predictive accuracy was achieved on PDXs not used to train the model: MCC=0.47 (PR=0.62 and RC=0.81). **(right)** *Cetuximab (1 fts)*: Using the same input data and evaluation protocol, the best single-gene marker of Cetuximab sensitivity was the mutational state of ACR. The RF-OMC model provided a slightly lower level of prediction than that of this standard single-gene procedure: MCC=0.59 (PR=0.90 and RC=0.56).

In the next three subsections, we further analyse the three predictors found to have the highest accuracies and hence are the most useful models to predict which forthcoming PDXs will be responsive.

### Predicting BRCA PDX response to Binimetinib

The best multi-gene predictor for BRCA PDXs treated with Binimetinib, a MEK1/2 inhibitor, was RF-OMC applied to GEX data comprising the expression values of 22,665 genes. Our analysis discarded the other three molecular profiles for Binimetinib-BRCA (SNV, CN and CNA), as they were substantially less predictive than the GEX profile in this case. RF-OMC practically offered the same performance as RF-all (MCC of 0.57 vs 0.56, respectively). However, RF-OMC identified 14 out of these 22,665 genes as the more informative to predict BRCA PDX response to Binimetinib. The resulting RF-OMC predictions are hence optimal combinations of the expression values of only these 14 genes, whereas RF-all was trained on all 22,665 GEX features.

Figure 3 displays the performance of this multi-gene predictor compared to that of the best single-gene marker for Binimetinib-BRCA (the mutational state of PABPC3 with P=0.02 from a two-sided Fisher’s exact test). The multi-gene predictor achieves a more substantial discrimination between sensitive and resistant markers than the PABPC3 marker. This is also indicated by a higher MCC (0.57 vs 0.24). To help to understand what MCC values represent in terms of achieved discrimination, we also indicated FP and FN errors from both classifiers.

### Predicting BRCA PDX response to Paclitaxel

The best multi-gene predictor for BRCA PDXs treated with Paclitaxel was RF-OMC applied to somatic mutation data comprising the presence or absence of a SNV in 15,232 genes. Our analysis discarded the other three molecular profiles for Paclitaxel-BRCA (GEX, CN and CNA), for being less predictive than the SNV profile in this case. The resulting RF-OMC model employs two out of these 15,232 genes (MUC20 and UPK3BL). RF-based combination of these two mutational states provide strongly better prediction than a RF model using the mutational states of all the genes (MCCs of 0.49 and -0.07, respectively).

Incidentally, we would like to highlight that RF-all constitutes an embedded feature selection technique^42^. RF-OMC performs generally better because it promotes a much more stringent feature selection, which has been found to be more suitable in most cases. We illustrate this point with paclitaxel-BRCA-SNV as an example. RF-OMC with only two SNV features achieves a MCC of 0.49. By contrast, with all features and the same data, RF-all obtains a far worse MCC of practically zero (MCC=-0.07). Out of these 15,232 features, only 3374 are actually used by any of the 1000 trees forming the RF-all model, as each RF tree only uses those features providing the best discrimination among the m_try_ randomly chosen at each node. The poor performance indicates that 3374 features are still far too many for this particular case, which is much better predicted by a RF model employing the two most predictive features.

Figure 4 visualises the high performance of this two-gene predictor on Paclitaxel-BRCA. The performance of the best single-gene marker, also shown, is very poor. This marker is the mutational state of HYDIN, a gene coding for Protein Phosphatase 1’s Regulatory Subunit 31. While we found this sensitising mutation to be the most strongly associated with the cytotoxic drug Paclitaxel in BRCA (P=0.04; two-sided Fisher’s exact test), its performance in left-out PDXs suggests that it is a spurious correlation.

### Predicting CRC PDX response to Cetuximab

The best multi-gene predictor for CRC PDXs treated with Cetuximab was RF-OMC applied to somatic mutation data. The other three molecular profiles for Cetuximab-CRC (GEX, CN and CNA), were discarded for being less predictive than the SNV profile in this case. We identified four out of these 15,232 genes whose combined mutational states provide better prediction than a RF model using the mutational states of all genes (MCCs of 0.47 and 0.39, respectively).

Figure 5 visualises the higher performance of this four-gene predictor on Cetuximab-CRC. The performance of the best single-gene marker, also shown, is even higher in this case. This marker is the mutational state of ACR (Acrosin), whose association to this targeted drug is the strongest across the 26 cases (P=0.0003; two-sided Fisher’s exact test). Unlike with RF-OMC, most prediction errors correspond to sensitive PDXs that were not correctly identified as such (FN=7 vs FP=1).

### Multi-gene predictors generally offer substantially higher recall than single-gene markers

In the three cases analysed in Figures 3-5, multi-gene predictors exhibit a substantially higher recall than the corresponding best single-gene markers. More concretely, recalls of 0.91, 0.88 and 0.81 (multi-gene) versus recalls of 0.68, 0.00 and 0.56 (single-gene). Figure 6 shows that this is actually a strong general trend: 23 out of 26 studied cases have a higher proportion of correctly predicted sensitive PDXs using the multi-gene markers. We recently observed a similar trend when using the standard version of RF to predict *in vitro* drug response from SNV data^12^, which is here confirmed *in vivo*.

**Figure 6:**
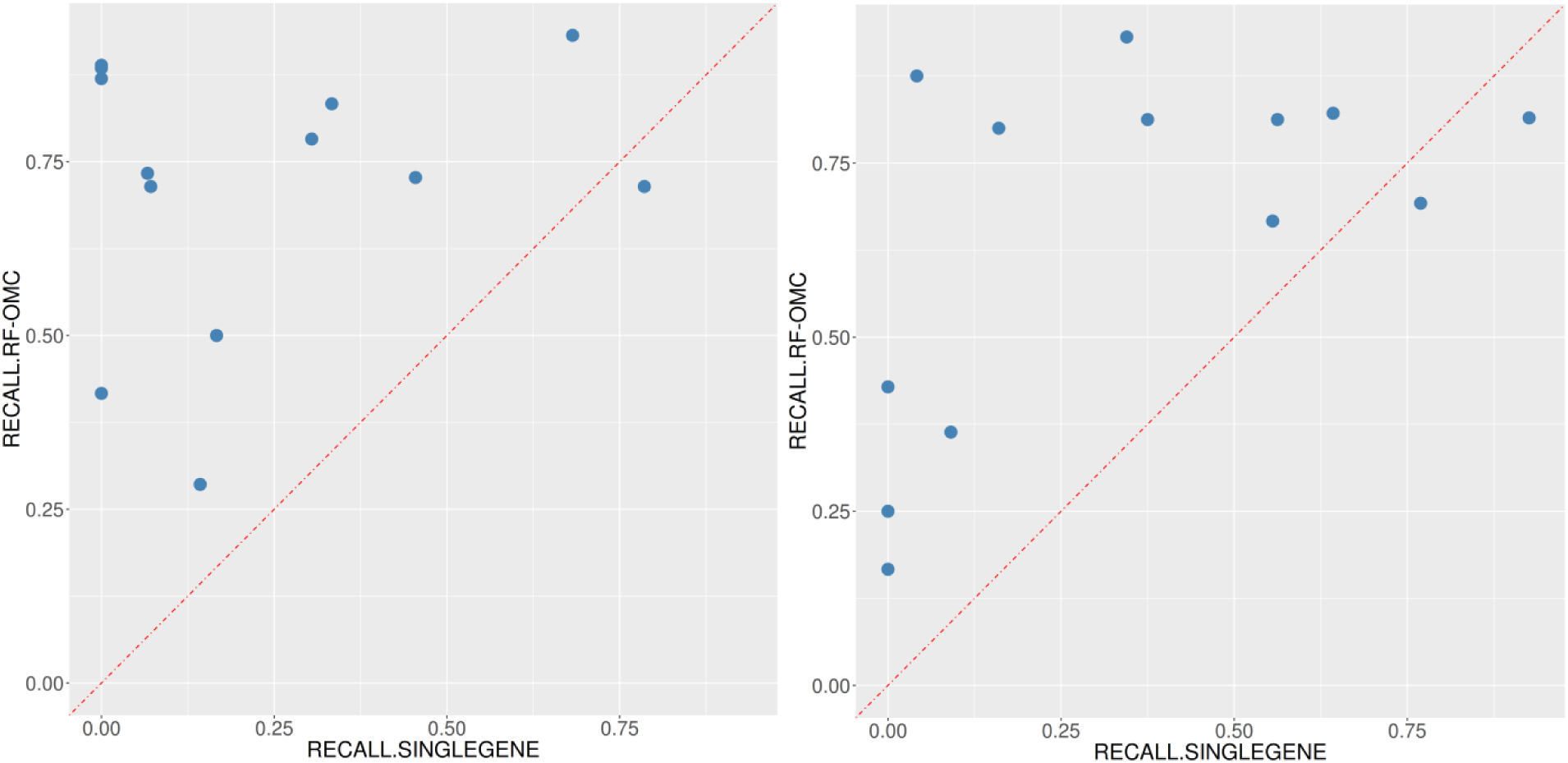
ML multi-gene classifiers exhibit a higher recall than single-gene markers in 23 of the 26 treatment-cancer type pairs. **(left)** In 12 of the 13 treatments for BRCA, multi-gene markers achieve a higher recall than the single-gene marker. The exception was the BGJ398-KIF20B association. **(right)** In 11 of the 13 treatments for CRC, multi-gene markers achieve a higher recall than the single-gene marker. These two single-gene markers with higher recall are the association BYL719-TMEM184A and the association BYL719+LJM716-LSR. Note that these three genes (KIF20B, TMEM184A and LSR) all have very high prevalence (47%, 61% and 85%, respectively) in addition to having the lowest p-value as sensitive marker in their respective cancer types.

In the three cases that do not follow this trend, a slightly higher proportion of correctly predicted sensitive PDXs was found using single-gene markers. The first case is BGJ398-BRCA and its best single-gene marker is the drug-gene association BGJ398-KIF20B. The other two cases are in CRC and have as best markers the following drug-gene associations: BYL719-TMEM184A and BYL719+LJM716-LSR.

## DISCUSSION

Combining multiple gene alterations via ML has resulted in better discrimination between sensitive and resistant PDXs in 19 of the 26 analysed cases. More importantly, this ML approach has determined which is the most predictive classifier type and molecular profile for each treatment – cancer type pair. Despite training on data from practically the same PDXs within a cancer type, we found that some treatments can be predicted much better than others (this is true for both cancer types). The results show that an effective way to improve the prediction of a given case is to evaluate several model types. For instance, the collective prediction of these 26 cases would have been much worse if we had only considered the standard RF algorithm (see for instance the difference between the MCCs of square and triangle signs for drug LKA136 in Figure 2). Likewise, the same undesirable outcome would have been occurred if only the GEX profile had been available (see MCCs of blue signs in Figure 2). This new knowledge is displayed in Figure 2 and fully reported in the Supplementary Tables.

We have also seen that even ML algorithms with built-in FS can often struggle to provide predictive classification models. To further mitigate the problem of overfitting caused by high-dimensional data^43^, we have introduced and evaluated RF-OMC. OMC ranks features by their individual ability to discriminate between resistant and sensitive tumours (e.g. via a t-test), with only the highest ranked being used to train the ML model. Thus, such univariate filters may miss co-operativity effects among features. In principle, wrapper FS techniques such as Recursive Feature Elimination (RFE)^40^ should improve model accuracy by capturing these effects. However, at least in the related problem of cancer prognosis prediction^41^, univariate filters generally outperform RFE despite the computational cost of RFE being much higher. In this same study, embedded (built-in) FS techniques such as LASSO did not generally outperform univariate filters either. Instead of using a fixed predetermined cutoff (e.g. the 100 top-ranked genes as in that study^41^), a novel aspect of OMC is that the complexity of the ML model is optimised for the drug, cancer type, molecular profile and available data. As the dimensionality of the employed data is optimally reduced for the considered case, thousands of less informative gene alterations are not included in model building. This is often an advantage, as the least informative of these features are probably irrelevant and hence harm classifier performance.

In practice, we have found that OMC complements the standard version of RF on this type of problems (16 of the 26 cases were better predicted by RF-OMC). More importantly, RF-OMC predicts 11 cases with an MCC of at least 0.3, whereas RF-all only predicts 4 cases with at least that accuracy. Furthermore, while RF-OMC predicts 21 of the 26 cases better than random, this is only the case for 16 of the 26 cases predicted by RF-all. Interestingly, unlike here with multi-omics features of tumours, FS did not generally result in more accurate ML models when using chemical features of drug molecules in the related problem of QSAR^44^. In our study, RF-all only outperforms RF-OMC once among the most predictive cases (i.e. those with an MCC of at least 0.3, see Supplementary Figure 6), when predicting BRCA PDX response to HDM201 based on SNV profiles. This probably due to limitation of this initial version of RF-OMC, where features are ranked without taking into account their possible synergies with the rest of features and the cut-off might be too tight. These exceptions also serve as a reminder of the importance to consider multiple algorithms.

As anticipating which classifiers will work best on a case is currently not possible, we benchmarked three algorithms building models of varying complexity. In this way, their relative performance can shed some light into the intrinsic non-observable complexity of each case. For instance, only a given list of genes could be controlling drug response for tumours of that type and that control could be preferentially exert at a given omics level. If a single-gene marker works best, it is likely that such subset is actually reduced down to that single gene and the omics level is SNV (e.g. the mutational status of ACR as a predictor of CRC PDX response to Cetuximab in Figure 2 right). However, it could also be that mutations in other genes influence drug response as well, but we do not have sufficient data to detect more complex cooperativity patterns. In cases where RF-OMC works best, a few genes are likely to be controlling drug response via the most predictive molecular profile (e.g. the mutational status of NR1H2, TLK2 and CTSA to predict BRCA PDX response to LKA136 in Figure 2 left), although again more data could result in more complex gene interactions being exploited and thus models with higher predictive accuracy. Lastly, in cases where RF-all obtains the best performance (e.g. an RF model trained on all SNV features to predict BRCA PDX response to HDM201 in Figure 2 left), this suggests that more features than those considered by the OMC strategy must be positively contributing to the prediction of drug response.

Indeed, an important advantage of RF-OMC over standard RF models is that only a few genes need to be profiled to predict whether a PDX is responsive or not. Concise gene lists in a highly predictive model are valuable for interpretation and clinical application purposes. Take for instance the 14 genes forming part of the Binimetinib-BRCA GEX predictor (Figure 3). RF-OMC unveiled that this gene list is a promising starting point for mechanistic studies. Such studies would aim at explaining how the nonlinear interplay between the expression values of these genes accurately predicts BRCA PDX response to Binimetinib. On the other hand, concise gene lists permit cheaper and faster clinical implementation. For example, instead of carrying out three whole-exome molecular profiles per tumour sample, we now know that it suffices to determine the mutational status of just two genes to predict BRCA tumour response to Paclitaxel (Figure 4). This example also highlights that the most responsive tumours to a cytotoxic drug can also be accurately predicted. The latter indicates that the applicability of precision medicine in current standard of care oncological therapeutic regimes is not restricted to targeted agents, but also includes cytotoxic chemotherapy^12,45^. While this is not the aim of the study, we have commented on the cancer relevance of the genes selected to predict treatment response for the three best RF-OMC predictors. This literature review can be found in the supplementary information file (pages 7-14).

We have also discovered that multi-gene predictors of *in vivo* drug response generally have higher recall than single-gene markers, which confirms *in vivo* previous findings *in vitro*^12,39^. The recall of a single-gene marker will be necessarily poor in all cases in which the prevalence of the mutation is much lower than the response rate. It is therefore not surprising that the three cases where markers have higher recall than RF-OMC in Figure 6 are based on genes with high to very high prevalence (KIF20B, TMEM184A and LSR with respective prevalence across tumours of 47%, 61% and 85%). Albeit exceptions, this general trend makes sense because a marker by construction can only detect those responsive tumours with the actionable mutation. In other words, the marker is blind to responsive tumours arising from alternative molecular mechanisms as illustrated by the example in Figure 1. By contrast, a ML algorithm can implicitly learn all such mechanisms from the data itself. Therefore, an important conclusion is that, without generating any additional data, many more patients should benefit from precision oncology by applying multiple ML algorithms to existing clinical pharmacogenomics data. Algorithms able to generate classifiers combining a few data-selected gene alterations such as RF-OMC are particularly promising.

Given the accuracy of some of these predictors, their eventual application to translational clinical pharmacogenomics research would be highly beneficial for the biomedical community and would ultimately improve clinical decision making. Likewise, our study has made a set of data modelling recommendations that can be applied to the analysis of any similar data set. As PDXs capture the diversity and complexity of their originating tumours^8^, the translational potential of our approach to the clinical setting is anticipated. Improved predictors for Paclitaxel and Cetuximab, both of which are standards of care for breast and colon cancers, respectively should impact on cancer treatment effectiveness, and consequently clinical practice in the near future. Our approach is also a useful tool to identify improved predictors for drug responses of compounds in drug development (e.g. Binimetinib), supporting the use of ML for patient selection in clinical trials. Beyond these immediate clinical applications, our study shows that ML can provide specific information to improve our understanding of cancer biology. Before drug testing, ML can identify those sensitive PDXs not harbouring any actionable mutation (these would have therefore been missed by existing single-gene markers). Going further, the OMC strategy provides a concise list of gene alterations that control drug response in the considered cancer type. An alternative hypothesis explaining drug reponse can be generated by combining both streams of information.

## METHODS

### NIBR-PDXE data

The NIBR-PDXE data set^29^ is publicly available as the Supplementary Table 1 at http://www.nature.com/nm/journal/v21/n11/full/nm.3954.html#supplementary-information. This Excel file has five tabs named RNASeq_fpkm, copy_number, pdxe_mut_and_cn2, PCT_raw_data and PCT_curve_metrics. The first three tabs contain three molecular profiles of the xenografted tumours. The RNASeq_fpkm tab contains gene expression values. The copy number tab contains the actual copy number of each gene. Copy number is also available as a categorical variable at the pdxe_mut_and_cn2 tab (this table also contains detected mutations per gene). Around 400 PDX models were profiled at each of these omic levels. The other two tabs are for treatment response data. The raw response data tab (PCT_raw_data) includes the percentage of tumour volume change relative to tumour volume at the start of treatment (%∆TVol) of each treated PDX recorded every 3-4 days. Lastly, the processed response data tab (PCT_curve_metrics) includes the categorisation of PDX responses into one of four classes calculated from raw response data. Further information about how this data set was generated can be found in the original study^29^.

### Processing treatment response data for modelling

For each treated PDX, we retrieved its category from the processed response data. We also calculated its category from raw response data as indicated by Gao et al.^29^. This calculation was based on the variables Best Response (the minimum value of %∆TVol for t ≥ 10 days) and Best Average Response (the minimum value of the set of average responses spanned by all t values with t ≥ 10 days). In particular, CR (Complete Response) was assigned if Best Response < -95% and Best Average Response <-40%; PR (Partial Response) if -95% ≤ Best Response < -50% and -40% ≤ Best Average Response <-20%; SD (Stable Disease) if -50% ≤ Best Response < 35% and -20% ≤ Best Average Response < 30%; otherwise PD (Progressive Disease) was assigned as the response category. Retrieved and calculated response categories differ in 277 of the 4758 PDX-treatment pairs. Although such discrepancies were small (mostly swaps between contiguous categories) and not numerous (5.8% of the cases), we decided to use the calculated categories in these cases so that all PDX-treatment pairs were categorised following the same set of rules. Gao et al.^29^ further subdivided PDX response into two classes: responders as those PDXs exhibited some level of sensitivity to the treatment (CR, PR or SD) and non-responders (PD) as those resistant to the treatment.

### Processing molecular profiling data for modelling

For each gene, the Single-Nucleotide Variant (SNV) feature for that gene is assigned a value of 1 if at least one SNV was detected in this gene region (i.e. reported in the pdxe_mut_and_cn2 tab). If no SNV was detected in this gene, the gene was labelled as Wild Type (WT) and the SNV feature was assigned a value of 0. This encoding scheme is commonly used^46–48^, as it has the advantage of leading to a much less sparse data instances vs features matrix than if a binary feature was defined for each SNV (a gene typically has several SNVs). In this way, each PDX profiled at the SNV level was characterised by a set of 15,232 binary features (one feature per gene). On the other hand, the copy_number tab contains the actual copy number (CN) of each gene as determined by Gao et al.^29^. This is the CN profile composed by 23,853 real-valued features (one per gene). Gao et al. categorised these measurements as follows: Amp5 if the gene is moderately amplified with copy number in the range >= 5 and < 8, Amp8 if the gene is strongly amplified with copy number >= 8 and Del0.8 if the gene is deleted with copy number <=0.8 (these per-gene CN categories were reported in pdxe_mut_and_cn2). As previously done elsewhere^46,47^, we binarised copy-number data to generate less sparse features: the Copy-Number Alteration (CNA) feature of a gene has a value of 1 for aberrant copy number (Amp8, Amp5 or Del0.8) and 0 otherwise. Thus, each PDX profiled at the CNA level is described by a set of 21,534 binary features (one per gene). By contrast, the fourth molecular profile, Gene EXpression (GEX), is directly the data provided in the RNASeq_fpkm tab. Thus, each PDX profiled at the GEX level is characterised by 22,665 real-valued features (one per gene as well).

### Processed data sets for modelling

Only a part of the PDX models from NIBR-PDXE have been both treatment-response and molecularly profiled. A previous cancer pharmacogenomics modelling study showed that it is possible to predict drug response of held-out tumours with a ML model trained on just 35 tumours^49^. As we are not aware of successful studies using smaller training sets, we focused on the two cancer types with the highest numbers of profiled PDXs per treatment, Breast Cancer (BRCA) and Colorectal Cancer (CRC), where all but one of these 26 treatment-cancer type pairs had at least 35 PDXs. Overall, a set of 13 treatments were administered to BRCA PDX models and another set of 13 treatments were administered to CRC PDX models. The first two tabs of Supplementary Table 1 state the numbers of sensitive and resistant PDXs per treatment for BRCA and CRC, respectively.

### Measuring the predictive performance of a classifier

The pharmacogenomics data set for a given cancer type, molecular profile and the i^th^ treatment can be represented as

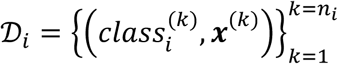

Where ***x*** is a high-dimensional vector with the features from the considered profile and the i^th^ treatment has been administered to n_i_ PDX models. In these binary classification problems, positive data instances are PDXs sensitive to the considered treatment (class=sensitive), whereas negatives are resistant PDXs (class=resistant). Note that, while they have slightly different meanings, we use the terms responder and sensitive PDX interchangeably as it is customary (same applies to the terms non-responder and resistant PDX).

Each of these data sets is employed to train classifiers to predict the class of a PDX from its corresponding molecular profile. Predictive performance is always reported on PDXs not used to train the classifier making the predictions. In particular, classifiers not employing model selection are evaluated with standard leave-one-out cross-validation (LOOCV). Moreover, classifiers employing model selection are evaluated with nested LOOCV to avoid overestimating their performance^50,51^. It is worth noting that nested LOOCV is nothing but a standard LOOCV where the model optimised (selected) with the training set of a given fold is applied to the test set of that fold (i.e. instead of training and testing the same model per fold).

Once observed classes are compared to predicted classes, PDXs in the considered data set can be broken down into true positives, true negatives, false positives and false negatives (their numbers being TP, TN, FP and FN, respectively). Thus, the discrimination offered by a classifier can be summarised by the Matthews Correlation Coefficient (MCC)^52^

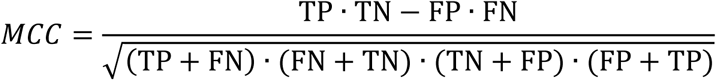

MCC can take values from -1 to 1, where 1 means that the classifier provides perfect agreement between observed and predicted classes, -1 indicates a perfect disagreement and 0 means that the classifier performance is equivalent to that of predicting the class at random.

To investigate how the two sources of error contribute to the overall predictive performance represented by MCC, we also calculate Precision (PR) and Recall (RC) for each predictor. PR and RC are two classical metrics^53^ whose definitions are:

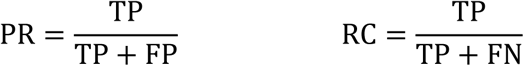

In this study, a PR value of 0 would mean that all the PDXs predicted sensitive by the classifier are actually resistant, whereas a PR value of 1 would indicate that all the PDXs predicted responsive were experimentally confirmed to be responsive. On the other hand, RC is 0 when none of the sensitive PDXs is correctly identified as such, whereas RC is 1 if no sensitive PDX is missed by the classifier.

### Multi-gene classifiers with built-in Feature Selection (FS)

Some ML algorithms can construct classifiers with built-in FS to mitigate the impact of the high dimensionality of data on their generalisation to unseen data. Random Forest (RF)^54^ is one of these algorithms, as it generates trees that ignore irrelevant features by construction and thus is often found effective in modelling high-dimensional omic data^55^. We used the recommended values for RF hyperparameters (1000 for the number of trees and the square root of the number of considered features for m_try_). We preferred this to tuning these hyperparameters for each training set, as RF tuning generally results in just marginal improvements at the cost of being much more computationally expensive^38,56,57^. As no model selection was carried out for this algorithm, standard LOOCV was performed to estimate the performance of RF using all the features (RF-all) on each data set (treatment-cancer type-molecular profile). To assess the variability introduced by the stochastic character of RF, we perform 10 repetitions of LOOCV per case, each using a different random seed (see corresponding boxplots in supplementary figures 4 and 5).

The proportion of responsive and non-responsive PDXs changes from case to case (see tabs data_by_treatment_BRCA and data_by_treatment_CRC in the Supplementary Tables). Although class imbalances are not strong, these could still introduce some loss in performance. Thus, we enabled class weighting in the RF algorithm (R package 'randomForest' version 4.6-12), which counterbalances class imbalances by putting a heavier penalty on misclassifying the minority class. The misclassification penalty of the minority class was set to the proportion of the majority class, thus promoting RF trees that are equally accurate regardless of the class.

### Multi-gene classifiers with Optimal Model Complexity (OMC)

An effective way to improve predictive performance is to reduce the dimensionality of the data. Here data dimensionality can be defined as the number of considered features over the number of PDXs. One route to reduce dimensionality is hence to use more training data, but these are usually not available. An alternative route is to only consider the most informative features in the data, thus typically discarding the many thousands of less informative features (hence strongly reducing data dimensionality while retaining most the initial information content).

However, the optimal number of features and their identities depend on various factors (treatment, profile, cancer type and data set). Consequently, we designed OMC as a strategy to build ML models employing only the most relevant features. In a nutshell, OMC is made of three modules: one to rank features according to their relevance to treatment response, another to train a ML model per considered subset of features and third one to select the optimal model among those trained. Regarding ranking features, we used p-values from two-sided Fisher’s exact tests to rank binary features within a given binary profile (SNV or CNA) and p-values from two-sided unpaired t-tests to rank real-valued features (GEX or CN). For each profile, treatment and cancer type, we considered n/2 subsets of features (n is the number of PDXs available for that case): the subset with the top 2 features, that with the top 3 features,…,that with top n/2 features and finally all features for that profile. This limit of n/2 ensures that the ML algorithm will not be challenged by high-dimensional data (i.e. all trained models will have at least two data points per considered feature), except for the run using all features intended to find out whether the case requires more than the top n/2 features. Lastly, the best among these n/2 models is selected as that with the highest LOOCV MCC. To estimate its performance, nested LOOCV MCC is calculated with model selection in the inner loop^50,51^ (the same values of the hyperparameters used for RF-all were also used here for RF-OMC). Sometimes the best RF-OMC model is that only employing the top 2 or the top 3 features (i.e. more complex models are not more predictive). As the RF model reduces in these cases to a set of redundant shallow trees, we are probably wasting computer time, as running RF with 10-50 trees should do just as well.

### Random model based on the prior probability of each case

The following protocol was followed for each case. First, the proportion of sensitive PDXs in each LOOCV training fold was taken as an estimate of the probability of a PDX to be sensitive. Second, a random number between 0 and 1 was generated to decide whether the held-out PDX in the LOOCV test fold was predicted sensitive or not according to the estimated prior probability. Third, the process was iterated over all LOOCV folds. Once the response class of every PDX had been predicted in this way, a single MCC value was computed from predicted and actual classes of the PDXs. Lastly, we repeated this process 10 times with the same 10 different random seeds employed with RF-OMC and RF-all. From the 10 MCCs for the case and the other 10 MCCs for the RF-model (either RF-OMC or RF-all), we carried out a one-sided paired t-test to determine whether the performance of the RF model was better than that of this random model (P<0.05). Results can be found in supplementary figures 2 to 5.

### Single-gene markers

We identified the best single-gene marker for each of the 26 treatment-cancer type pairs using exactly the same data as RF-all and RF-OMC via LOOCV. The SNV profile was used as the source of detected somatic mutations, as there is currently a strong interest in using them as pharmacogenomic markers in oncology^5^. For each fold, the response of the PDX held-out in the test fold was predicted with the most significant sensitive marker (i.e. predicted sensitive if a SNV is detected in the marker gene, predicted resistant otherwise). Such marker was determined by calculating two-sided Fisher’s exact tests across training fold PDXs, one per gene, leading to p-values and effect sizes (ϕ^58^) for 15,232 genes. The gene with the lowest p-value among those constituting sensitive mutations (ϕ>0) was identified. The operation was repeated for each fold resulting in a LOOCV predicted class for each treated PDX and thus the LOOCV MCC for the best marker. After evaluating its predictive performance, we recalculated each best single-gene marker using now all the data so that these markers are ready to be used on forthcoming tumours. These results are in the Supplementary Tables (single-gene_marker tab).

## Supporting information

## ABBREVIATIONS

FN: number of False Negatives
FP: number of False Positives
PDX: Patient-Derived tumour Xenograft model
NIBR-PDXE: Novartis Institutes for Biomedical Research - PDX encyclopedia
MCC: Matthews Correlation Coefficient
PR: PRecision
RC: ReCall
SG: Single-gene RF: Random Forest
OMC: Optimal Model Complexity
TN: number of True Negatives
TP: number of True Positives
WT: Wild-Type
LOOCV: Leave-One-Out Cross-Validation
CN: Copy Number
CNA: Copy-Number Alteration
SNV: Single-Nucleotide Variant GEX: Gene EXpression

## SUPPLEMENTARY INFORMATION

**Supplementary Tables:** Description of processed NIBR-PDXE data and the discovered multi-omics predictors of *in vivo* treatment response.

**Supplementary Figures:** Visualisation of the criteria for categorising PDX responses, p-values of ML models with respect to a null distribution, comparison RF-OMC vs RF-all across cases and plots showing how considering multiple classifiers and molecular profiles collectively improve predictive accuracy.

## AUTHOR CONTRIBUTIONS

P.J.B. conceived the study and designed the experiments. P.J.B wrote the manuscript with the assistance of all authors. L.N. carried out the numerical experiments. All authors analysed the results and contributed to their discussion.

## ACKNOWLEDGMENTS

This work has been carried out thanks to the support of the 911 Programme PhD scholarship from Vietnam National International Development (L.N.), IPC PhD scholarship (A.B.), Horizon 2020 work program of the Marie Curie Actions (G.G.), Canceropôle PACA (S.N.) and INSERM (P.J.B).

## CONFLICTS OF INTEREST

The authors declare no conflicts of interest.

## DATA AVAILABILITY

This study did not generate any data.

