## Supplementary material for "Machine learning models to predict *in vivo* drug response via optimal dimensionality reduction of tumour molecular profiles"

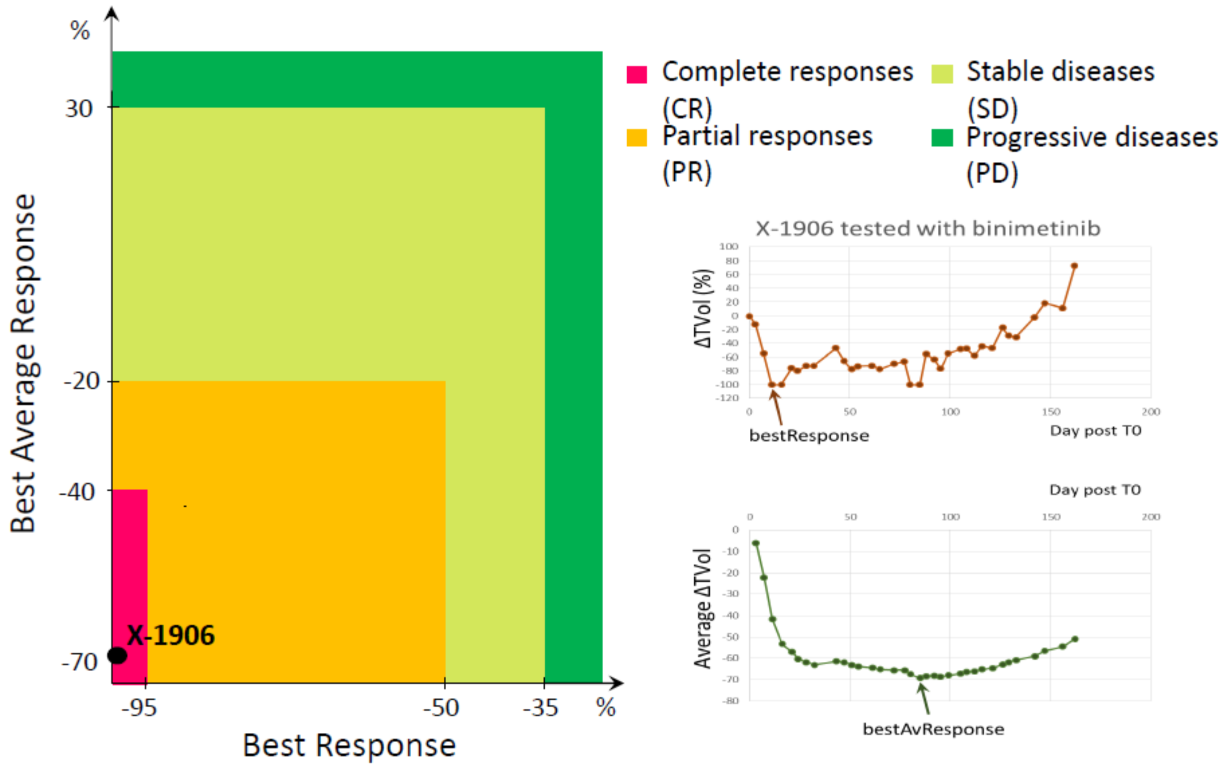


**Supplementary Figure 1. Visualising criteria employed by the NIBR-PDXE to categorise PDX responses to a treatment from the temporal evolution of tumour volume grow.** NIBR-PDXE employed a set rules, which and rules are visualised here for clarification purposes, including an example of a PDX with complete remission (X-1906). Both rules and nomenclature above are defined in the Methods section of the paper (e.g. ΔTVol corresponds to the % of variation in tumour volume with respect to initial tumour volume).

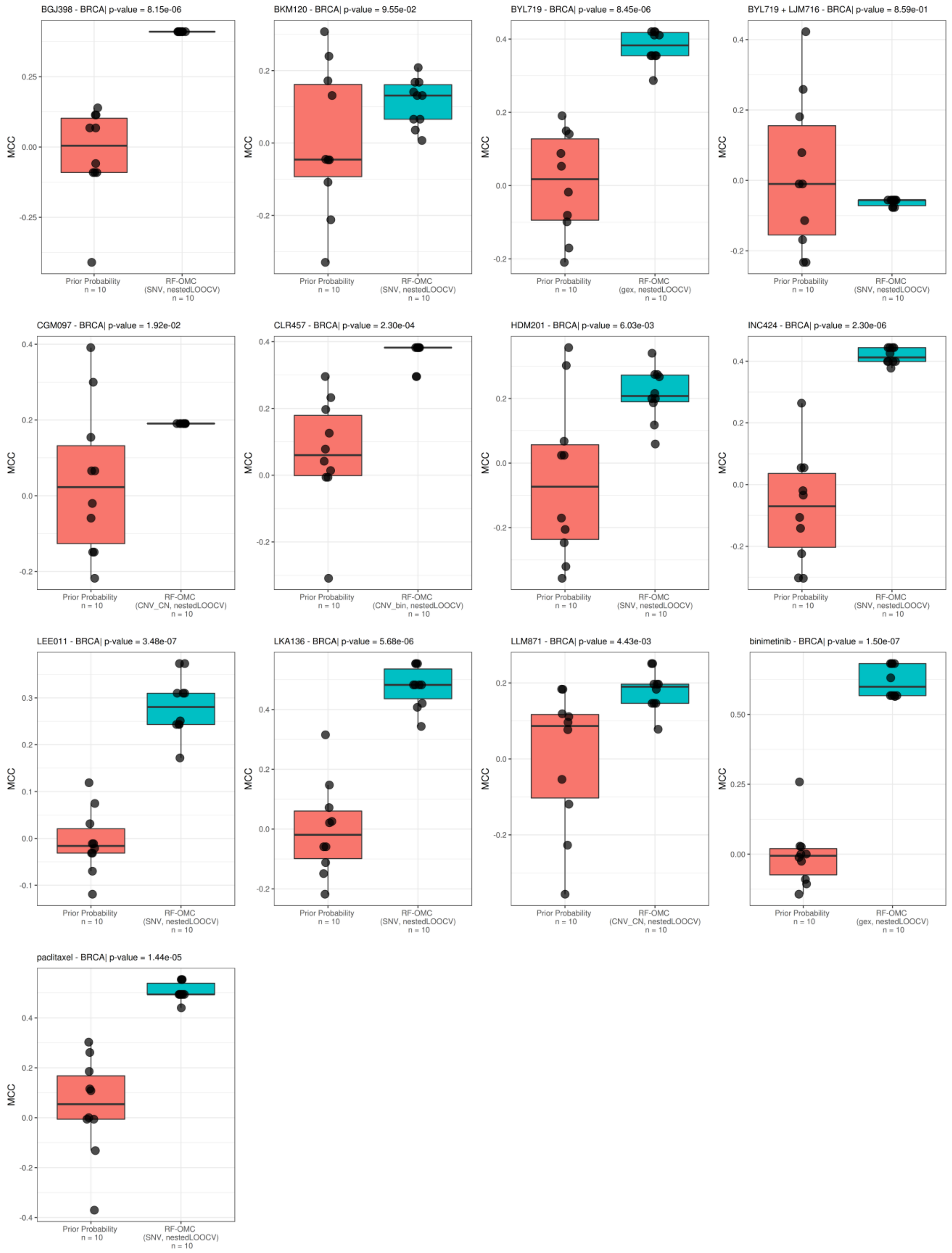


**Supplementary Figure 2. RF-OMC versus random predictions for each of the 13 treatments administered to Breast Cancer (BRCA) PDX models.** The random model is based on the prior probabilities for the case (see Methods section). For each of the 13 treatments, a magenta boxplot summarises the MCCs of the 10 runs of RF-OMC. The turquoise boxplot contains the MCCs of the 10 runs of the random model using prior probabilities. Each calculated MCC is presented as black point. From the p-values reported on the title of each plot, we can see that RF-OMC predicts 11 of the 13 treatments significantly better than random (p-value < 0.05; one-sided paired t-test).

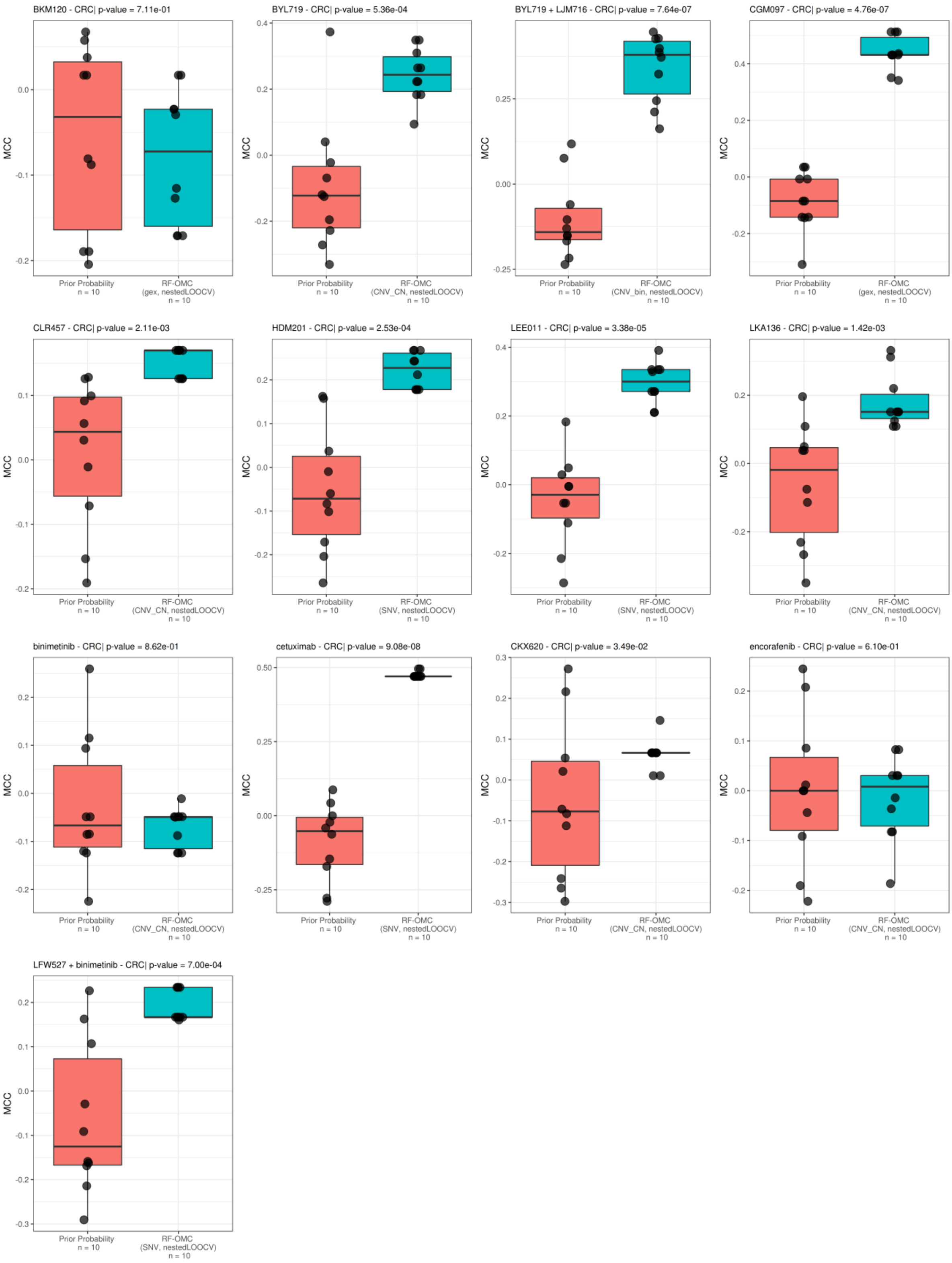


**Supplementary Figure 3. RF-OMC versus random predictions for each of the 13 treatments administered to Colorectal Cancer (CRC) PDX models.** The random model is based on the prior probabilities for the case (see Methods section). For each of the 13 treatments, a magenta boxplot summarises the MCCs of the 10 runs of RF-OMC. The turquoise boxplot contains the MCCs of the 10 runs of the random model using prior probabilities. Each calculated MCC is presented as black point. From the p-values reported on the title of each plot, we can see that RF-OMC predicts 10 of the 13 treatments significantly better than random (p-value < 0.05; one-sided paired t-test).

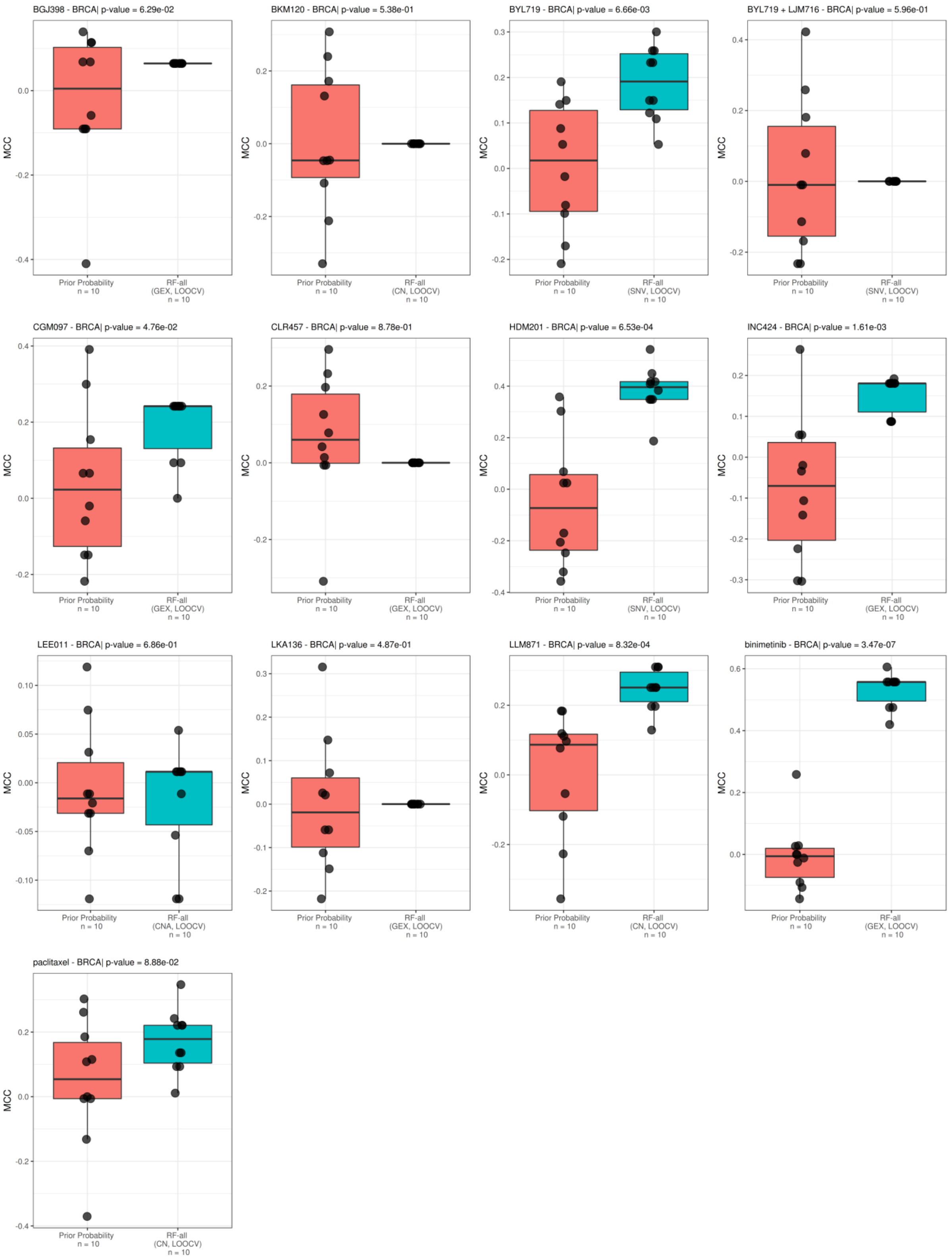


**Supplementary Figure 4. RF-all versus random predictions for each of the 13 treatments administered to BRCA PDX models.** The random model is based on the prior probabilities for the case (see Methods section). For each of the 13 treatments, a magenta boxplot summarises the MCCs of the 10 runs of RF-all. The turquoise boxplot contains the MCCs of the 10 runs of the random model using prior probabilities. Each calculated MCC is presented as black point. From the p-values reported on the title of each plot, we can see that RF-all predicts 6 of the 13 treatments significantly better than random (p-value < 0.05; one-sided paired t-test).

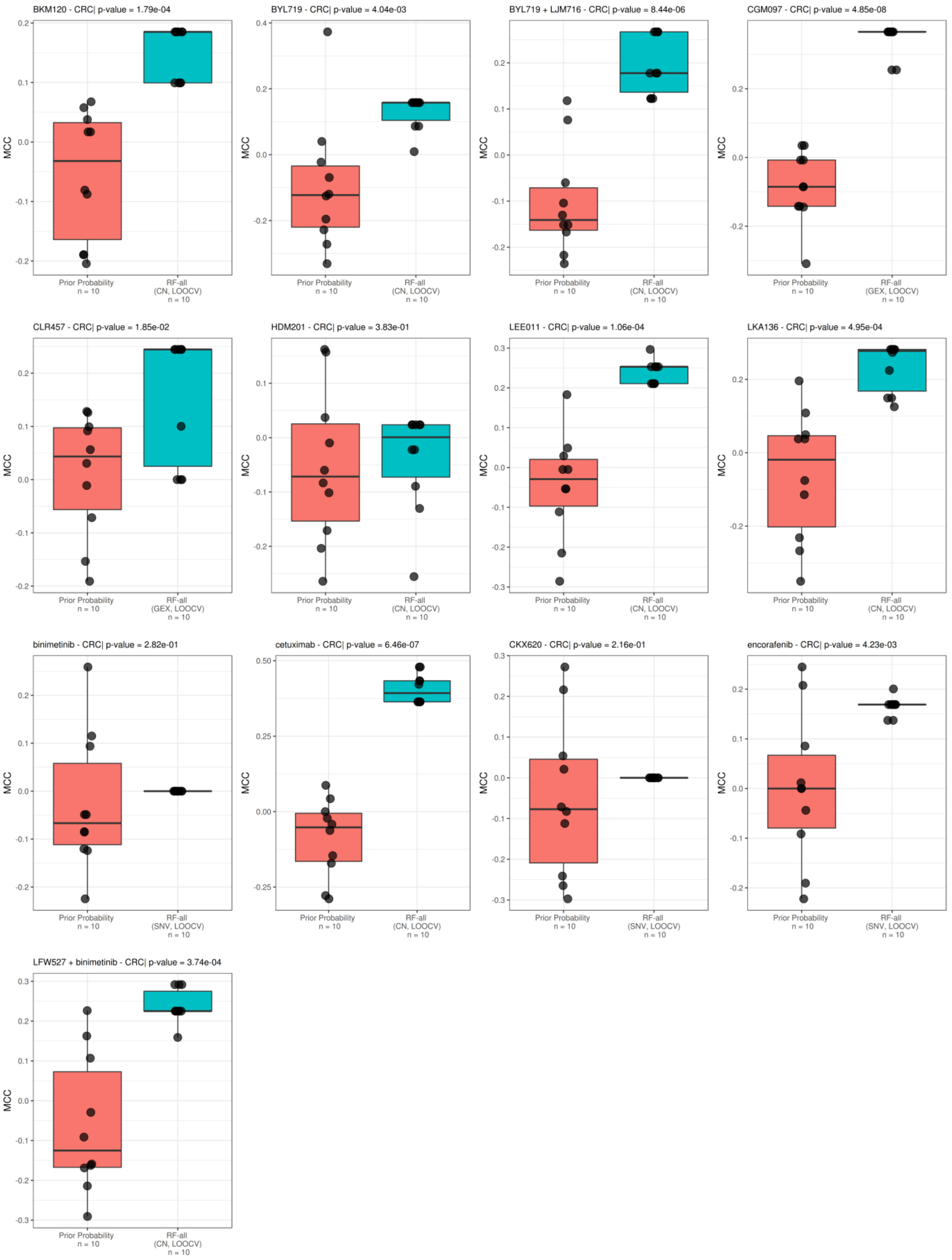


**Supplementary Figure 5. RF-all versus random predictions for each of the 13 treatments administered to CRC PDX models.** The random model is based on the prior probabilities for the case (see Methods section). For each of the 13 treatments, a magenta boxplot summarises the MCCs of the 10 runs of RF-all. The turquoise boxplot contains the MCCs of the 10 runs of the random model using prior probabilities. Each calculated MCC is presented as black point. From the p-values reported on the title of each plot, we can see that RF-all predicts 10 of the 13 treatments significantly better than random (p-value < 0.05; one-sided paired t-test).

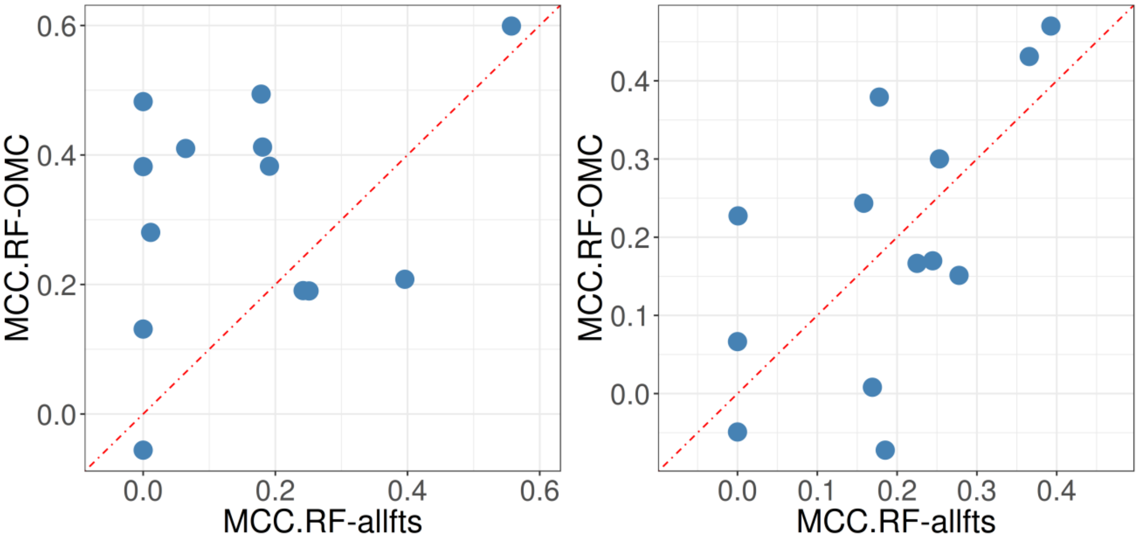


**Supplementary Figure 6.** Performance comparison of RF-OMC and RF-all models in predicting tumour response. **(left)** BRCA RF-OMC models achieved better MCC than RF-all models in 9 of 13 treatments tested on BRCA PDXs (7 RF-OMC models with MCC of at least 0.3, for only 2 RF-all models). **(right)** RF-OMC models achieved better MCC than RF-all models in only 7 of 13 treatments tested on CRC PDXs (4 RF-OMC models with MCC of at least 0.3, for only 2 RF-all models).

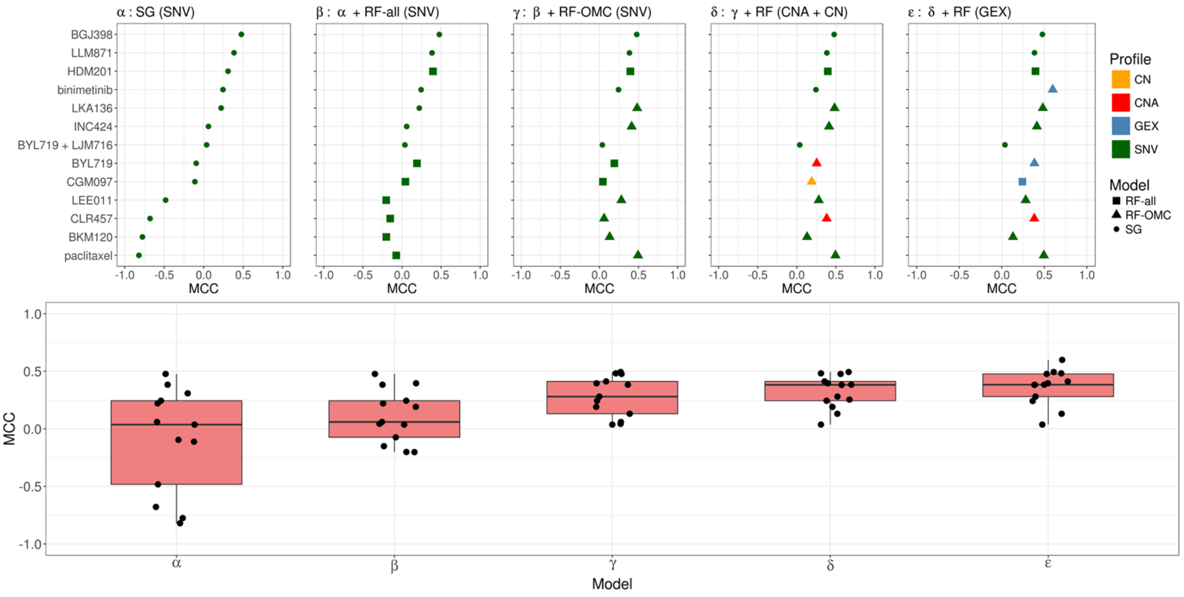


**Supplementary Figure 7. Prediction performance improves as more classifier and profile types are considered for predicting response of BRCA PDXs to 13 treatments. (Top)** In each of these five plots (α to **ε**), each row shows the cross-validated MCC of the best predictor for that treatment as we consider more classifiers and tumour profiles. **α** – Single-gene (SG) markers using Single-Nucleotide Variant (SNV) data. **β** – Also considering Random Forest model with all features (RF-all) leads to an improvement in prediction performance in 7 out of 13 treatments. Both models (SG and RF-all) considered SNV data. **γ** – also considering RF-OMC helped increase the accuracy in predicting 6 out of 13 treatments in comparison to plot β. All three models (SG, RF-all and RF-OMC) were trained using SNV data. **δ** – Models considering both SNV and copy-number data, including real-valued (CN) and binary copy-number alterations (CNA). The addition of these profiles results in an increase in performance in 3 out of 13 treatments in comparison to plot γ. **ε** – Models also considering the gene expression (GEX) profiles, the performance is improved in 3 out of 13 treatments compared to plot δ. With all classifiers and available profiles considered, 10 of the 13 treatments can now be predicted with MCC>0.25. By contrast, in the common case of using a single classifier and profile represented by α and using the same PDXs, only 3 treatments can be predicted with this level of accuracy. (**Bottom**) Each boxplot summarises the MCC values of the employed classifiers across the 13 treatments for each case mentioned above.

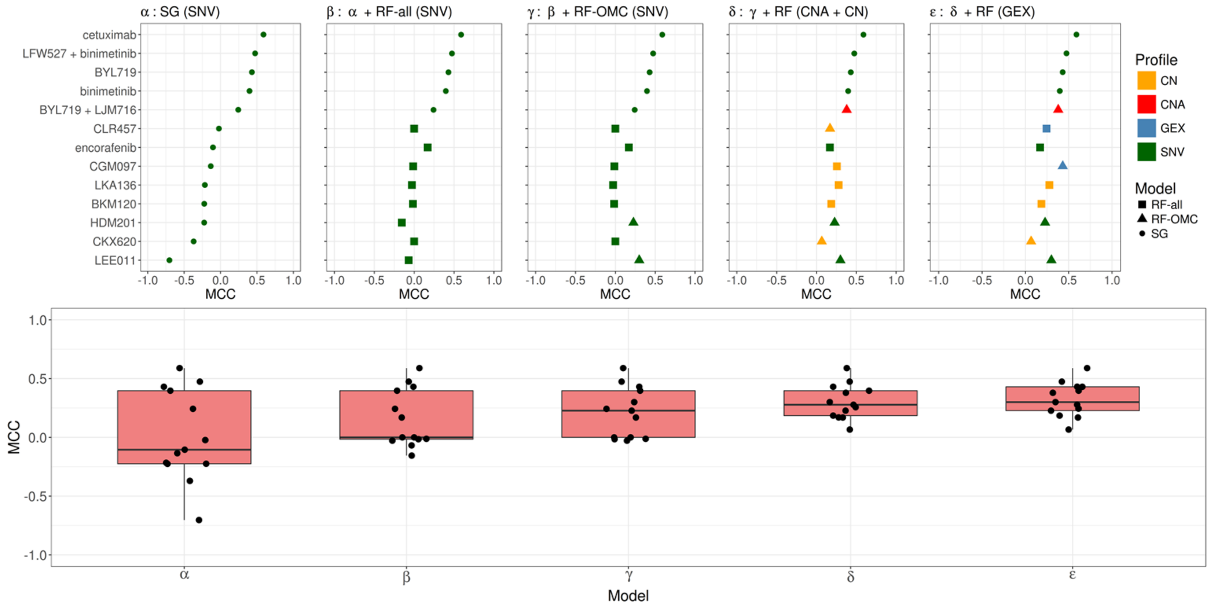


**Supplementary Figure 8. Prediction performance improves as more classifier and profile types are considered for predicting response of CRC PDXs to 13 treatments. (Top)** In each of these five plots (α to **ε**), each row shows the cross-validated MCC of the best predictor for that treatment as we consider more classifiers and tumour profiles. **α** – Single-gene (SG) markers using Single-Nucleotide Variant (SNV) data. **β** – Also considering Random Forest model with all features (RF-all) leads to an improvement in prediction performance in 8 out of 13 treatments. Both models (SG and RF-all) considered SNV data. **γ** – also considering RF-OMC helped increase the accuracy in predicting 2 out of 13 treatments in comparison to plot β. All three models (SG, RF-all and RF-OMC) were trained using SNV data. **δ** – Models considering both SNV and copy-number data, including real-valued (CN) and binary copy-number alterations (CNA). The addition of these profiles results in an increase in performance in 6 out of 13 treatments in comparison to plot γ. **ε** – Models also considering the gene expression (GEX) profiles, the performance is improved in 2 out of 13 treatments compared to plot δ. With all classifiers and available profiles considered, 8 of the 13 treatments can now be predicted with MCC>0.25. By contrast, in the common case of using a single classifier and profile represented by α and using the same PDXs, only 4 treatments can be predicted with this level of accuracy. (**Bottom**) Each boxplot summarises the MCC values of the employed classifiers across the 13 treatments for each case mentioned above.

### RF-OMC classifier to predict the response of BRCA PDXs to paclitaxel

This model exclusively employs the mutational status of two genes MUC20 and UPK3BL. The MUC20 gene encodes for a member of the mucin protein family. Mucins are high molecular-weight extracellular glycoproteins which are secreted by epithelial cells to form an insoluble mucous barrier. Many mucins have been found to be abnormally expressed or glycosylated in adenocarcinomas as well as associated with carcinogenesis, tumour invasion and poor patient outcome (1). In particular, the MUC20 gene has also been found to promote aggressive phenotypes in ovarian cancer (2), and to be the predictor of recurrence and poor outcome in

CRC (3). High MUC20 expression is associated with response to chemotherapy in esophageal squamous cell carcinoma (4). The UPK3BL gene encodes for a protein in the uroplakins protein family. It has been found among the up-regulated genes in rhabdoid glioblastoma tumour, suggesting that this gene may be functionally involved in this type of cancer (5).

**RF-OMC classifier to predict the response of BRCA PDXs to binimetinib**

This model uses the expression levels of just 14 genes: CRB3, NDUFA1, MPG, ECI1, ING2, KIF9, TSTD1, FAM100A, TCEAL3, HAGH, PEX11G, SNORA72, SNORA70 and PIN1. The CRB3 gene encodes for a protein in the CRB protein family, which plays various roles in the control of cytokinesis, ciliogenesis and the formation of tight junctions between cells, as well as involves in the establishment of cell polarity in epithelial cells (6). Dysregulation of cell polarity proteins could play an important role in cancer development (6). The overexpression of CRB3 gene has been show to inhibit breast cancer cell growth and promote apoptosis *in vitro*, as well as to reduce tumour growth *in vivo* (7). Meanwhile, a reduced expression of CRB3 was proved to induce carcinogenesis in mouse kidney epithelial cells (8). Another study has shown that CRB3 affects the expression of the epithelial-mesenchymal transition (EMT) transcriptional repressor (9). The EMT process – the biological process in which the polarized epithelium undergoes changes in the cell cytoskeleton, loses epithelial features and acquires mesenchymal characteristics like migration and invasion – is known to play essential roles in cancer development (10). The MPG gene encodes for an DNA repair enzyme, *N*-methylpurine-DNA glycosylase. This gene has increased expression in breast cancer cell lines in comparison to normal breast epithelial cells (11). The ING2 gene is a member of the inhibitor of growth (ING) protein family. These proteins function in DNA repair and apoptosis. The expression of ING2 gene has been found significantly reduced in human melanoma, and the under-expression of ING2 could be an important event in the initiation of melanoma development (12). Other studies have suggested that ING2 gene could be an tumour supressor gene in the carcinogenesis of head and neck squamous cell cancer (13) and in lung cancer (14). Although no evidence has been found for the KIF9 gene relating to carcinogenesis or cancer progression, this gene has been shown to be required for chromosome alignment and mitotic progression (15). The TCEAL3 gene encodes for a protein in the transcription elongation factor A family, which may function as a nuclear phosphoprotein modulating transcription in a promoter-dependent manner. A study on the association between gene expression and drug resistance in colorectal cancer patients discovered that TCEAL3 is one of the genes that significantly associate with patient survival (16). The HAGH gene encode for the enzyme glycoxylase II in charge of catalyzing the glutathione-dependent metabolism of cytotoxic methylglyoxal, which protects cells against cellular damage and apoptosis (17). An increasing body of evidence has suggested that glycoxylase II participates in the initiation and progression of urological maglinancies, including prostate cancer (18,19), renal cancer (20,21) and bladder cancer (22). PEX11G is the gene encoding for the Peroxisomal Biogenesis Factor 11 Gamma – a protein that joins in the regulation of number and sizes of the peroxisomes in the cells. Although no direct association has been found between the PEX11G gene and carcinogenesis, a study pointed out that this gene is one of the direct target of TP53 (23). SNORA70 and SNORA72 are two members of the small nucleolar RNA genes. While the roles of these two genes have not been comprehensively studied, certain other members of this gene family were found to act as oncogenes or tumour suppressors genes *in vitro* (24,25). Last but not least, the gene PIN1 encodes for the Peptidyl-prolyl cis/trans isomerases, which catalyzes the cis/trans isomerization of peptidyl-prolyl peptide bonds. PIN1 plays an important role in breast development, as well as it is a target of several oncogenic pathways and was found overexpressed in breast cancer (26,27). Besides, the overexpression of PIN1 also promotes *in vitro* cell growth of osteosarcoma cell lines (28).

**RF-OMC classifier to predict the response of CRC PDXs to cetuximab**

This model considered the gene expression levels of only four genes: ACR, DENND4B, NOTCH1 and RPL22. Interestingly, this model considers the gene NOTCH1, whose association with cancer is well known. This gene encodes a transmembrane receptor participating in the Notch signaling pathway. Aberrations in the genes of this pathway could have a large impact on cellular division leading to cancer (29). This pathway has been found dysregulated in CRC, where the upregulation of NOTCH1 is associated with poor survival outcome (30). Although no mutation of this gene has been reported in CRC, a study found that NOTCH1 amplification in metastatic CRC patients could lead to worse survival in this subgroup of patients (31). The gene RPL22 encodes for a cytoplasmic ribosomal protein that is a component of the 60S subunit. Mutations and abnormal expression of RPL22 gene has been reported in various types of cancers, including T-cell acute lymphoblastic leukemia (32), endometrial cancer (33), colorectal cancer (34), non-small cell lung cancer (35) and gastric cancer(36). We could not find any study linking ACR or DENND4B to CRC, but the high predictive accuracy of this classifier demonstrates the important role of these genes. Experimental studies intended to unravel the role of these genes is therefore promising.

Among the genes resulted from our multigene RF-OMC classifiers, a high number of them have reported to be associated with cancer in the literature. It is noteworthy that these genes were selected via a data-driven approach. Therefore, these results suggested that the models found by RF-OMC are biologically relevant and can be further investigated for the application in practice.

2. Chen C-H, Shyu M-K, Wang S-W, Chou C-H, Huang M-J, Lin T-C, et al. MUC20 promotes aggressive phenotypes of epithelial ovarian cancer cells via activation of the integrin

β1 pathway. Gynecol Oncol [Internet]. 2016 Jan [cited 2018 Jun 27];140(1):131–7. Available from: http://www.ncbi.nlm.nih.gov/pubmed/26616226

http://www.ncbi.nlm.nih.gov/pubmed/21052543

36. Nagarajan N, Bertrand D, Hillmer AM, Zang Z, Yao F, Jacques P-É, et al. Whole-

genome reconstruction and mutational signatures in gastric cancer. Genome Biol [Internet]. 2012 Dec 13 [cited 2018 Nov 2];13(12):R115. Available from: http://www.ncbi.nlm.nih.gov/pubmed/2323766
